# Conditional targeting of phosphatidylserine decarboxylase to lipid droplets

**DOI:** 10.1101/2020.10.22.350470

**Authors:** Santosh Kumar, Chandramohan Chitraju, Robert Farese, Tobias Walther, Christopher G. Burd

## Abstract

Phosphatidylethanolamine is an abundant component of most cellular membranes whose physical and chemical properties modulate multiple aspects of organelle membrane dynamics. An evolutionarily ancient mechanism for producing phosphatidylethanolamine is to decarboxylate phosphatidylserine and the enzyme catalyzing this reaction, phosphatidylserine decarboxylase, localizes to the inner membrane of the mitochondrion. We characterize a second form of phosphatidylserine decarboxylase, termed PISD-LD, that is generated by alternative splicing of PISD pre-mRNA and localizes to lipid droplets and to mitochondria. Sub-cellular targeting is controlled by a common segment of PISD-LD that is distinct from the catalytic domain and is regulated by nutritional state. Growth conditions that promote neutral lipid storage in lipid droplets favors targeting to lipid droplets, while targeting to mitochondria is favored by conditions that promote consumption of lipid droplets. Depletion of both forms of phosphatidylserine decarboxylase impairs triacylglycerol synthesis when cells are challenged with free fatty acid, indicating a crucial role phosphatidylserine decarboxylase in neutral lipid storage. The results reveal a previously unappreciated role for phosphatidylserine decarboxylase in lipid droplet biogenesis.

## Introduction

Phosphatidylserine (PS) and phosphatidylethanolamine (PE) are structurally related aminophospholipids that fulfill structural roles as components of organelle membranes, provide specific biochemical and cellular functions in a variety of processes such as apoptosis and autophagy, and serve as metabolic precursors to other lipids (Calzada et al., 2016; Vance, 2008). PE is widely distributed throughout most organelles of the cell, though it is an especially abundant component of the mitochondrion. In contrast, PS is enriched in the cytoplasmic leaflet of the plasma membrane (PM) and, to a lesser extent, to membranes of organelles derived chiefly from the PM (i.e., endosomes) and the *trans*-Golgi network (Fairn et al., 2011). The distinct localizations of PE and PS within the cell are conserved throughout eukaryotic evolution suggesting that these lipids fulfill specific functions where they reside within the cell.

Phosphatidylethanolamine is produced by multiple pathways, including by decarboxylation of PS by phosphatidylserine decarboxylase (PSD) (Calzada et al., 2016; Vance, 2008), an enzyme that localizes to the inner membrane of the mitochondrion in human cells (Borkenhagen et al., 1961; Dennis and Kennedy, 1972; Kuchler et al., 1986; Percy et al., 1983; Schuiki and Daum, 2009; Zborowski et al., 1983). Mitochondrial PSD produces most, if not all, of the PE contained within mitochondrial membranes (Aaltonen et al., 2016; Shiao et al., 1995), as well as PE that is exported from the mitochondrion and distributed to other organelles via inter-organelle vesicular trafficking. However, the genomes of many organisms encode multiple PSD enzymes that localize to organelles of the late secretory and endosomal system (Di Bartolomeo et al., 2017). Interestingly, a yeast (*Saccharomyces cerevisiae*) phosphatidylserine decarboxylase has been shown to be targeted conditionally to the mitochondrion or to the endoplasmic reticulum depending on cellular metabolism; whereas growth on glucose favors mitochondrion targeting, growth on a fermentable carbon source favors localization to the endoplasmic reticulum (Friedman et al., 2018). These observations highlight organelle-specific targeting of PSD enzymes and establish that targeting of yeast Psd1 can be regulated by nutritional state.

In vertebrate organisms, PSD activity is encoded by a single gene, called *PISD* (Kuge et al., 1991). The annotated *PISD* gene product contains a canonical amino-terminal cleaved mitochondrion targeting signal, a membrane spanning segment, and a site of autocatalytic proteolysis that generates the α and β subunits from a single precursor (Chacinska et al., 2009; Clancey et al., 1993; Di Bartolomeo et al., 2017; Horvath et al., 2012; Kuge et al., 1991; Li and Dowhan, 1988; Trotter et al., 1993). Disruption of the PISD locus in the mouse genome results in embryonic lethality and fragmentation of mitochondria in viable cells derived from *Pisd*^−/-^ embryos (Kuge et al., 1991; Steenbergen et al., 2005; Tasseva et al., 2013). Similarly, yeast cells deleted of *PSD1* (*psd1Δ*) (Trotter et al., 1993), which encodes the major yeast PSD activity, also have fragmented mitochondria and are deficient in respiratory activity (Birner et al., 2001; Chan and McQuibban, 2012), indicating that PSD activity fulfills a conserved function(s) that is required for mitochondrial activity. In this study, we discovered a previously unrecognized form of PSD that localizes to the surface of the lipid droplet (LD), a storage organelle for neutral lipids, such as triacylglycerol and cholesterol esters. Depletion of both PSD isoforms by RNAi compromises triacylglycerol (TAG) storage in cells grown in high lipid conditions, revealing that PISD activity is critical for neutral lipid storage.

## Results and Discussion

### An alternatively spliced variant of PISD localizes to the lipid droplet

Although the human genome contains a single locus encoding PSD (called *PISD*), the genomes of many other eukaryotic organisms contain multiple genes encoding PSD enzymes that localize to other cellular compartments (Di Bartolomeo et al., 2017). This prompted us to closely examine the human proteome for evidence of additional PSD enzymes. A search of the National Center for Biotechnology Information (NCBI) human protein database using the protein sequence of human mitochondrial PISD (CCDS: 87016.1; RefSeq: NP_001313340.1) identified an uncharacterized form of PISD (CCDS: 13899.1; RefSeq: NM_014338.3) that differs from the query sequence at the N-terminus of the protein (Fig. 1A). The results of polymerase chain reaction (PCR)-based tests employing combinations of oligonucleotides derived from sequences that are unique to each transcript and a common downstream primer confirmed that both PISD transcripts are expressed in HeLa cells (Fig. 1B). Other organisms, chiefly fungi and plants, express multiple PSD proteins that are encoded by distinct genes (Di Bartolomeo et al., 2017). However, examination of the human and mouse PISD loci revealed the presence of exons found in both PISD transcripts, hence, the human and mouse genomes encode two forms of PISD that arise from the same locus by alternative pre-mRNA splicing. An analysis of all genomes in the NCBI database revealed that this unique form of PISD first appeared during evolution in fishes and is now present in all vertebrates examined. For reasons that are evident in the next section, we termed the ‘canonical’ mitochondrial PISD isoform “PISD-M” to indicate that it localizes exclusively to the mitochondrion, and the second isoform of PISD,;PISD-LD”.

**Figure 1:**
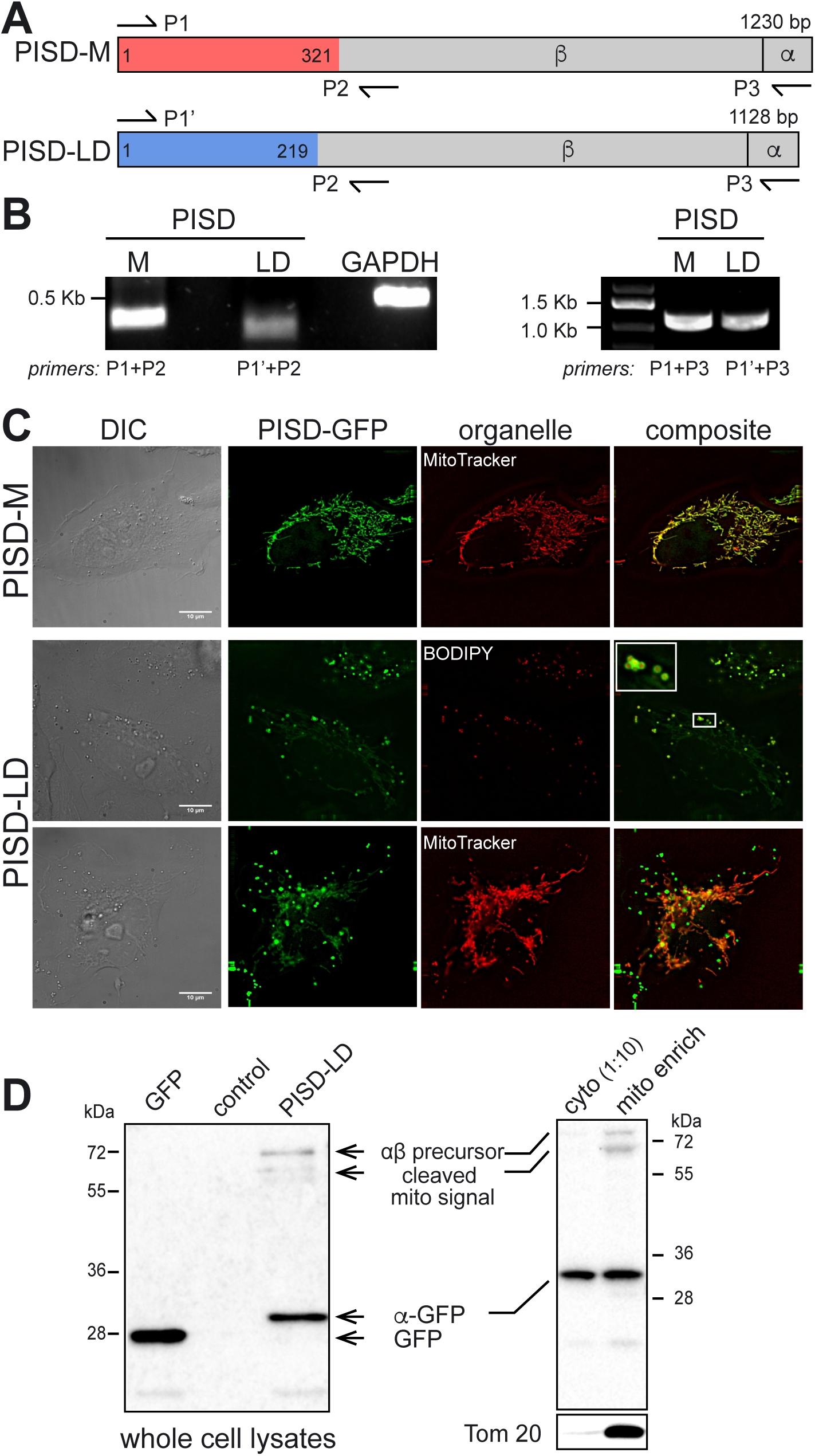
An alternatively spliced isoform of PISD localizes to the lipid droplet and mitochondrion. **(A)** Schematic representation of the two isoforms of PISD, PISD-M (mitochondrial form;1230 bp ORF) and PISD-LD (lipid droplet form; 1128 bp ORF). The unique segments of PISD-M and PISD-LD colored red and blue, respectively, and the region colored grey is common to both common forms. The position of auto-proteolysis, which generates the N-terminal *β* and C-terminal *α* subunits, is indicated by a vertical line. The arrows show the positions of PCR primers (e.g., P1, P2, etc) that were used to amplify segments of PISD-M and PISD-LD by RT-PCR. **(B)** Both PISD-M and PISD-LD isoforms are expressed in Hela cells. The indicated primer pairs were used to amplify the unique N-terminal segments of PISD-M and PISD-LD from cDNA synthesized from total cellular RNA. Amplification of GAPDH cDNA served as a control. The gel on the left shows results of amplification with primers unique to each mRNA and a common primer just downstream of the junction. The gel on the right shows results of amplification of each full length ORF. **(C)** Localization of GFP-tagged PISD-M and PISD-LD. Hela cells expressing PISD-M-GFP or PISD-LD-GFP were labeled with BODIPY red to visualize lipid droplets, or MitoTracker Red to identify mitochondria. The inset in the PISD-LD panel shows lipid droplet cores decorated on the surface with PISD-LD-GFP. Cells were grown in DMEM + 10% FBS. Images are single focal plane from a Z series. The scale bars represent 10 μm. **(D)** PISD-LD-GFP is proteolytically processed in HeLa cells and a small proportion of it co-purifies with mitochondria. Lysates of HeLa cells grown in DMEM + 10% FBS expressing GFP or PISD-LD-GFP were subjected to anti-GFP immunoblotting. The “GFP” lane contains lysate from cells expressing GFP alone, the “control” lane contains lysate from untransfected cells, and the “PISD-LD” lane contains lysate from cells expressing PISD-LD-GFP. The position of the GFP-tagged α subunit, which is ∼ 4 kDa larger than free GFP, is indicated. The positions of other, deduced PISD-LD-GFP precursors are indicated. On the right is shown an immunoblot of a mitochondrion-enriched subcellular fraction and a 1:10 dilution of cytosol. The blot was stripped and re-probed with antibody to Tom20, a mitochondrial protein (below).

The first 107 amino acids of PISD-M are unique to this isoform and this region contains a sequence that is predicted with high confidence (using MitoProt II; (Claros and Vincens, 1996)) to constitute a canonical cleaved mitochondrion targeting signal (Fig. 1A). Mitochondrial localization of PISD was first described on the basis of PSD enzymatic activity displayed by sub-cellular fractions (Dennis and Kennedy, 1972), but to our knowledge, direct visualization of the PISD protein in mammalian cells has not been reported. In our laboratory, commercially available antisera against PISD do not decorate mitochondria, so we constructed a PISD-M-GFP fusion gene, where GFP was fused to the C-terminus of the protein so as not to mask the mitochondrion targeting signal. We note that a yeast Psd1-GFP fusion protein complements a *psd1Δ* mutant (Friedman et al., 2018). As expected, PISD-M-GFP co-localized precisely with mitotracker, a dye that accumulates within the mitochondrion (Fig. 1C).

The first 73 amino acids of PISD-LD are unique to this form, but this region does not contain a predicted mitochondrion targeting signal. To determine subcellular localization of PISD-LD, we examined a C-terminally GFP-tagged form of PISD-LD (Fig. 1C). In cells grown in nutrient replete medium, PISD-LD-GFP decorated the surface of a variable number of small (< 1μm) round organelles and there was a faint tubular pattern reminiscent of the mitochondrial network. The cores of all of the punctate organelles decorated by PISD-LD-GFP stained with BODIPY dye, identifying them as lipid droplets (LDs) (Fig. 1C). No co-localization was found with various proteins of endosomes or lysosomes, which can also appear as puncta by fluorescence microscopy. We conclude that PISD-LD-GFP localizes in part to lipid droplets. Importantly, recent proteomics-based inventories of LD proteins in multiple cell types (U2OS, THP-1, SUM159) identified native PISD in purified lipid droplet fractions (Bersuker et al., 2018; Mejhert et al., 2020).

The faint tubular network decorated by PISD-LD-GFP stained with MitoTracker dyes (Fig. 1C), indicating that PISD-LD-GFP also localizes to mitochondria. This was confirmed with biochemical fractionation experiments where, for cells grown in nutrient-replete conditions, ∼15% of PISD-LD-GFP co-fractionates with a native mitochondrial protein, Tom20 (Fig. 1D). Importantly, anti-GFP immunoblotting of lysates of cells expressing PISD-LD-GFP or PISD-M-GFP shows that the major GFP-tagged species migrates at 32 kDa, corresponding in size to that of the alpha subunit-GFP fusion protein. Thus, neither the unique N-terminal segment of PISD-LD, nor GFP fused at the C-terminus, interferes with autoproteolytic processing reaction that generates the α and β subunits. In addition, a rare ∼80 kDa species, corresponding to the predicted size of the full-length precursor, and a smaller precursor form of ∼65 kDa, were observed. Based on the characterized processing intermediates of yeast mitochondrial PSD (Psd1) (Horvath et al., 2012), the 65 kDa species likely corresponds to a precursor that has been imported into the mitochondrion and its N-terminal mitochondrion targeting segment has been cleaved, but the autoproteolytic cleavage has not yet occurred. Consistent with this, both putative precursor species are enriched in a mitochondrion-enriched cell fraction, while the GFP-tagged α subunit is present in both the cytosol and mitochondria fractions (Fig. 1D).

### PISD-LD possesses overlapping LD and mitochondrion targeting signals

Because PISD-LD and PISD-M possess unique N-terminal sequences and localize distinctly within the cell, we hypothesized that the unique regions of each isoform confer distinct organelle targeting. Consistent with this, a protein composed of just the unique segment of PISD-M (amino acids 1-107) fused to the amino terminus of GFP localized to mitochondria (Fig. 2). Unexpectedly, a similarly constructed fusion protein containing the unique segment of PISD-LD (amino acids 1-73) did not localize to LDs; instead, it localized exclusively to mitochondria (Fig. 2). In addition, a fusion protein containing the segment that is common to PISD-LD and PISD-M (amino acids 74-375 of PISD-LD) also localizes to mitochondria suggesting that there are at least two mitochondrial targeting signals in human PISD isoforms.

**Figure 2:**
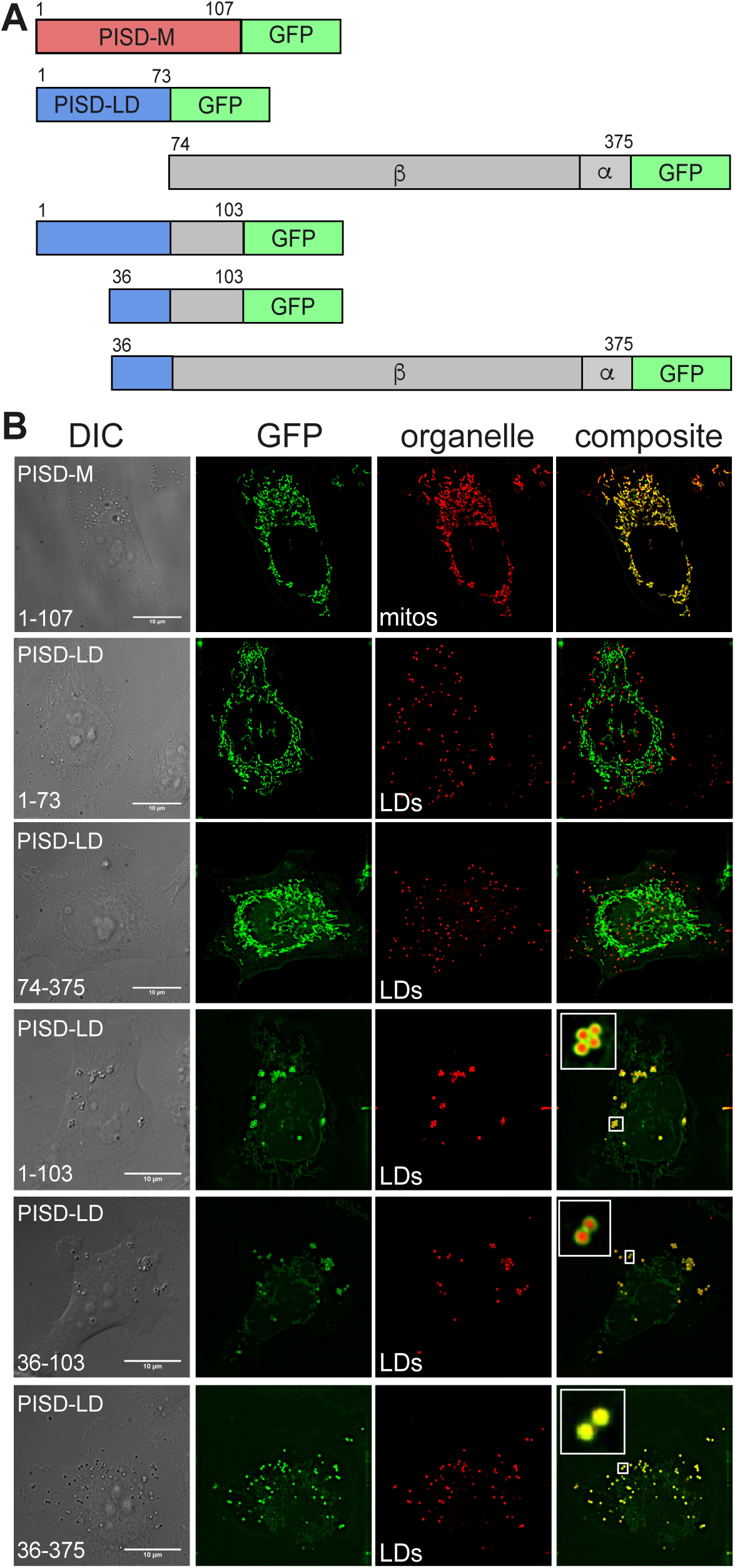
Identification of the lipid droplet targeting signal of PISD-LD-GFP. **(A)** Diagram of PISD-GFP fusion proteins. The indicated segments of PISD-M or PISD-LD were fused to the N-terminus of GFP. The numbers indicate the position of the first or last amino acid of the segment. **(B)** Micrographs of Hela cells expressing the indicated fusion proteins. Mitochondria were identified with MitoTracker Red and LDs were identified with BODIPY. A single focal plane from the approximate center of the cells is shown. Insets: PISD-LD-eGFP truncated form (1-103, 36-103 and 36-375 amino acids) localized to lipid droplets. The scale bars represent 10 µm.

The localization data indicate that a portion of PISD-LD that is common to both the mitochondrion and LD forms is required for LD targeting. To identify this LD targeting determinant, we determined localization of additional fusion proteins whose endpoints were selected on the basis of secondary structure predictions and hydropathy analysis (described below). Ultimately, these experiments revealed that a segment comprising amino acids 36–103 of PISD-LD is necessary and sufficient to confer LD targeting (Fig. 2). We note that this segment (fused to GFP) also localizes weakly to mitochondria, suggesting that LD and mitochondrion targeting are conferred by the same, or overlapping, sequences of the protein.

Several types of structural features of LD proteins have been identified that confer LD targeting (Kory et al., 2016). Some amphipathic alpha helices confer reversible targeting to the LD surface from the cytosol/aqueous phase, and a second type of LD targeting determinant consists of a hydrophobic segment that forms a lipid-embedded hairpin (Mejhert et al., 2020). Proteins containing the latter type include integral membrane proteins that target from the bulk endoplasmic reticulum (ER) to the surface of ER-associated LDs. Hydropathy analysis and secondary structural predictions of the PISD-LD sequence revealed the presence of a modestly hydrophobic segment (amino acids 47-63) and a putative amphipathic alpha helix (amino acids 70-93) within the minimal LD targeting determinants of PISD-LD.

The contribution of each feature to LD targeting was assessed by determining localization of mutant forms of PISD-LD-GFP (Figs. 2, 3). The length of the hydrophobic segment (15 amino acids) is not sufficient to span a typical biological membrane bilayer, leading us to postulate that it may constitute a membrane-embedded hairpin, similar to hydrophobic regions of other LD-associated proteins. This hypothesis was tested by reducing the hydrophobicity of this segment by changing three Leucine residues within it to Arginine (L_(51, 54, 55)_R). These changes resulted in a protein that fails to localize to the LD or to the mitochondrion (Fig. 3B), indicating that the hydrophobic character of this region is indeed required for proper targeting. Curiously, the hydrophobic segment of PISD-LD contains three Pro residues (Fig. 3A), resembling the “proline knot” motif that targets plant oleosin proteins to the LD (Abell et al., 2004; Chapman et al., 2012). To investigate a potential requirement for the proline residues in LD targeting, all three of them were changed to Leucine (P_(47, 56, 62)_L) and intracellular localization was determined. These substitutions still supported LD targeting, but also conferred targeting to ER tubules and large ER-associated puncta that do not stain with BODIPY, suggesting that they are aggregates within the ER membrane (Fig. 3B). In addition, although LD targeting was preserved by these Pro-to-Leu mutations, its expression resulted in fewer and larger LDs, raising the possibility that this mutant form of PISD-LD interferes with LD biogenesis. These data indicate that the hydrophobic, proline-rich segment of PISD-LD plays a critical role in targeting to both the LD and the mitochondrion.

**Figure 3:**
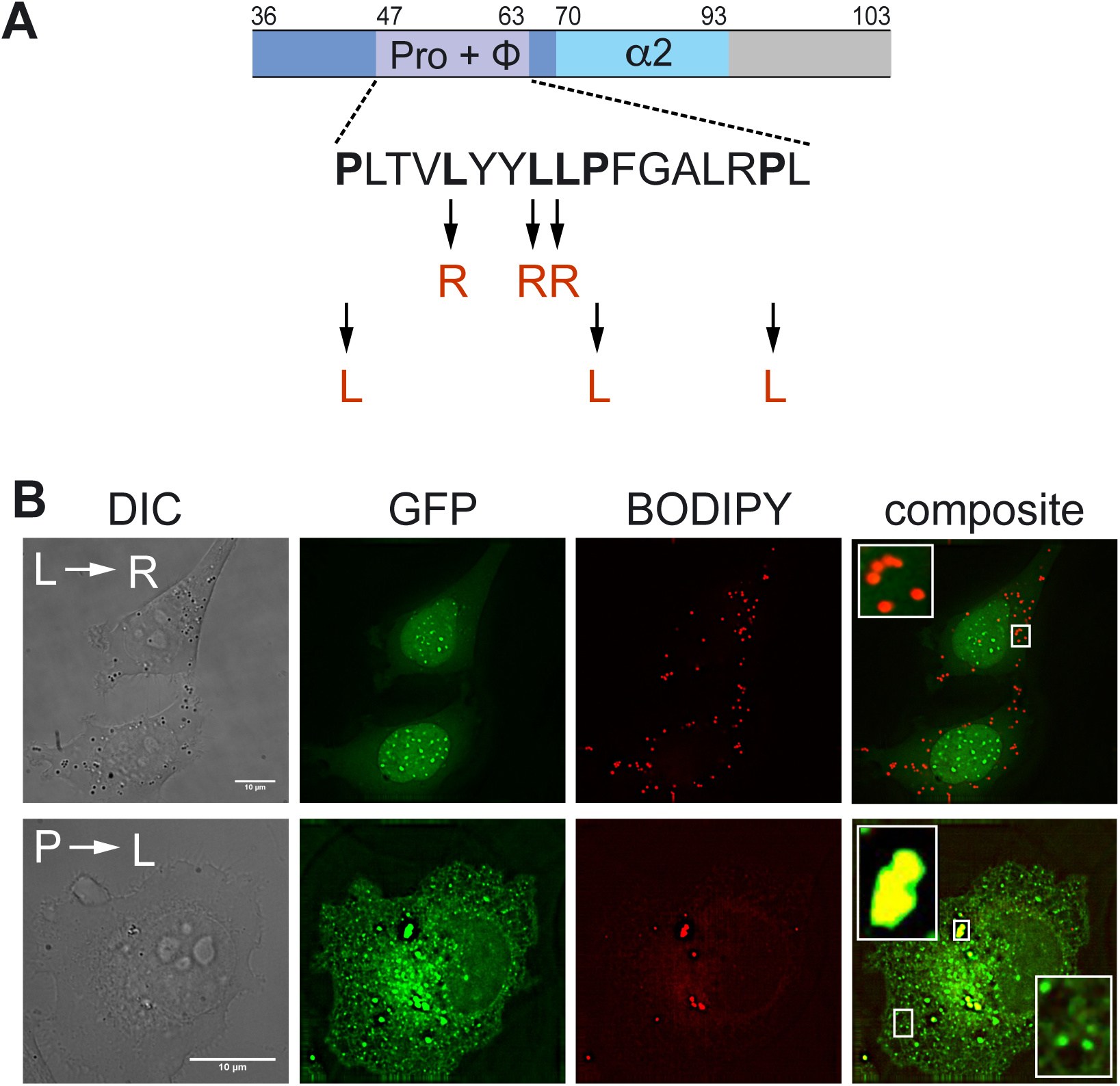
Hydrophobic region is required for proper targeting of PISD-LD to LDs. **(A)** Schematic representation of predicted structural features within the minimal LD targeting segment of PISD-LD-GFP. The sequence of the segment spanning amino acids 47-63, denoted “Pro + F”, is shown; it is rich in proline and hydrophobic resides. Two mutant versions of full-length PISD-LD-GFP fusion proteins were constructed. In one, the three indicated leucine residues (bold) were changed to arginine (denoted “L→R”), and in the other mutant, the three indicated proline residues (bold) were changed to leucine (denoted “P→L”). The segment spanning amino acids 70-93, denoted “α2”, is predicted to constitute an amphiphathic alpha helix. **(B)** HeLa cells expressing PISD-LD-GFP proteins. HeLa cells were transfected and stained with BODIPY. The insets show a higher magnification view of LDs in transfected cells. Images are single focal plane from a Z series. The scale bars represent 10 μm.

To address the role of the putative amphipathic alpha helix (amino acids 73-93) in PISD-LD targeting, we first assessed the role of residues comprising the hydrophobic face of the helix by mutagenesis (Fig. 4). Three Leu residues were changed to positively charged residues (Leu80Lys, Lue89Lys, Leu90Arg), thereby reducing the mean hydrophobicity of this segment from 0.42 to −0.03 (Fig. 4). These mutations result in a protein that does not localize to the LD, however, mitochondrion targeting is unaffected (Fig. 4B). A complementary mutant, in which two proline residues on the hydrophobic face on the helix were changed to Leu residues (Pro(74, 85)Leu), thereby increasing the mean hydrophobicity from 0.42 to 0.53 (Gautier et al., 2008)), results in a protein that localizes solely to LDs, with no mitochondrion localization apparent (Fig. 4B). These results indicate that the putative amphipathic alpha helix contributes to both LD and mitochondrion targeting.

**Figure 4:**
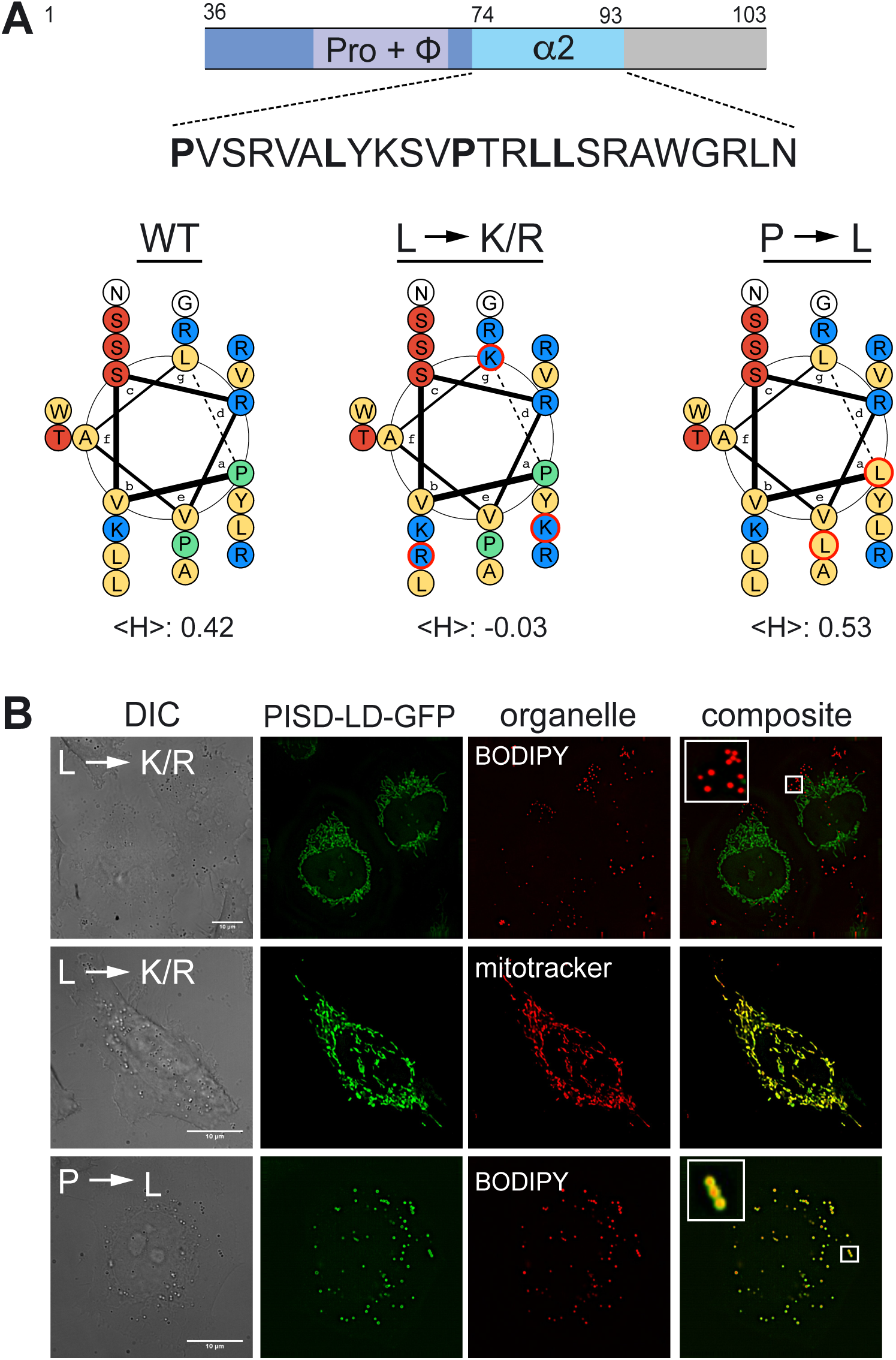
Amphipathic alpha helix is required for targeting of PISD-LD to lipid droplets. **(A)** Schematic representation of predicted structural features within the minimal LD targeting segment of PISD-LD-GFP. The sequence of the segment spanning amino acids 70-93 (denoted “α2”), predicted to form an amphipathic alpha helix, is shown. The bold leucine residues were changed to Lysine or Arginine (denoted “L→K/R”), and the bold Proline residues were changed to Leucine (denoted “P→L”). Helical wheel diagrams (amino acids 70-87) of PISD-LD of native sequence (“WT”) and the two mutants are shown with the calculated hydrophobicity of this region (“<H>“) is indicated below. Lysine and Arginine residues are colored blue, hydrophobic residues are colored yellow, Serine and Threonine residues are colored red, and Proline is colored green. **(B)** Localization analysis of PISD-LD mutants. Images are single focal planes from the approximate center of z series. Lipid droplets were identified by staining with BODIPY and mitochondria were identified by staining with MitoTracker. The scale bars represent 10 µm.

### PISD-LD targeting is regulated by nutritional state

As the lipid droplet is a neutral lipid storage organelle, localization of PISD-LD to both LDs and mitochondria raises the question, are these competing targeting outcomes that are controlled by cellular lipid metabolism? To address this, we monitored PISD-LD-GFP localization in cells grown in conditions that promote neutral lipid storage and expansion of LDs, or conditions that promote consumption and shrinkage of LDs. To promote LD expansion, the growth medium was supplemented with free fatty acid (oleate, 100 μM for 24 hours), and this resulted in the expected appearance of numerous clusters of enlarged LDs that are decorated by PISD-LD-GFP (Fig. 5A). In this population of cells, PISD-LD-GFP was found solely on LDs in 78% of cells and on both LDs and mitochondria in the remaining cells (Fig. 5B). Next, the medium of the oleate loaded cells was replaced with lipid-free medium to force consumption of existing LDs. As expected, after 24 hours in lipid-free medium, clusters of large LDs were no longer observed, and only small, isolated LD remnants were present. The proportion of cells in which PISD-LD solely decorated LDs was reduced to 10%, and the proportion of cells displaying localization to both LDs and mitochondria was increased to 87%. After 48 hours in lipid free medium, PISD-LD-GFP did not localize solely to LDs in any cell, and the proportion of cells displaying exclusive localization to mitochondria was increased to 68%. In the remainder of the population (32%), PISD-LD localized to both mitochondria and LD remnants. These results indicate that abundant free fatty acid in the growth medium favors targeting of PISD-LD-GFP to the surface of LDs, and that conditions that favor consumption/shrinkage of LDs result in targeting of PISD-LD-GFP to mitochondria.

**Figure 5:**
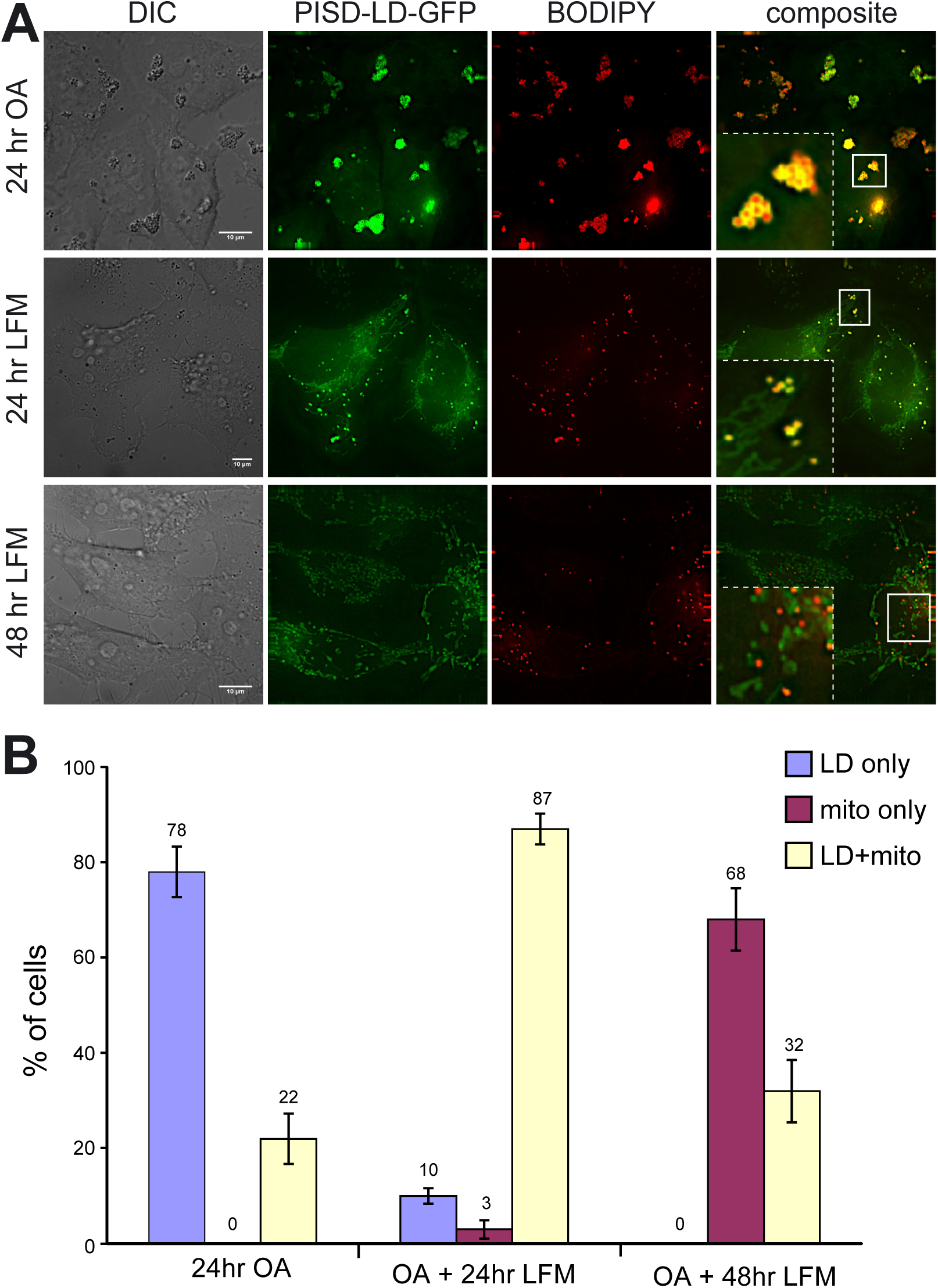
Fatty acid metabolism regulates PISD-LD-GFP localization. **(A)** Micrographs of cells expressing PISD-LD-GFP. Cells were transfected and maintained in DMEM + 10% FBS, then incubated for the indicated number of hours (prior to acquiring images) in medium containing oleic acid (“OA”; 100 μM) or in lipid-free medium (“LFM”), and then stained with BODIPY red to label lipid droplets. The scale bars represent 10 μm. **(B)** Proportions of cells (minimum of 60 cells analyzed per condition) with PISD-LD-GFP localized to lipid droplets and no mitochondria localization detected (“LD only”), localized solely to mitochondria and no lipid droplet localization detected (“mito only”), or to both lipid droplets and mitochondria (“LD+mito”). The means (indicated) and s.d. of three biological replicate experiments are plotted.

An interesting question for future studies regards the mechanism by which the LD targeting sequence of PISD-LD conditionally targets the enzyme to the LD or to the mitochondrion. Because we identified mutations in PISD-LD that result in strict localization to the LD or to the mitochondrion regardless of nutritional status, conditional targeting of the native enzyme reflects the outcome of competing processes. A similar situation was recently described for mammalian MLX-family transcription factors where LD expansion favors recruitment to the LD surface by binding of an amphipathic alpha helix located outside of the MLX basic helix-loop-helix domain, but MLX is released from the LD surface and traffics to the nucleus upon LD consumption. We suggest that conditional targeting of human PISD-LD-GFP is mediated by ‘sub-optimal’ targeting features, similar to dual targeting of yeast Psd1 between the mitochondrion and the ER, in which the efficiency of mitochondrion targeting is postulated to be reduced when cell growth is dependent on respiration (Friedman et al., 2018). For the case of human PISD-LD, a ‘weak’ mitochondrial targeting signal(s) in PISD-LD kinetically favors residence in the cytoplasm, thereby promoting residence on the LD surface, provided that there is sufficient surface area to accommodate PISD-LD. However, when LD surface area is limiting, the ‘weak’ nature of the LD targeting sequence of PISD-LD is insufficient to drive association with the crowded LD surface, thereby favoring targeting to the mitochondrion. We speculate that interactions between the hydrophobic segment and the amphipathic alpha helix are required for LD targeting, and perhaps with other parts of the enzyme, expose or mask LD targeting determinants.

### PISD promotes triglyceride storage

Similar to PISD-LD, CTP:phosphocholine cytidylyltransferase (CCT) is recruited to the surface of LDs in conditions favoring LD expansion, where it is activated to produce PC to maintain the LD glycerophospholipid monolayer (Krahmer et al., 2011). In cells depleted of CCT and challenged with oleate loading, enlarged LDs accumulate and triacylglycerol (TAG) content is increased (Krahmer et al., 2011). To address a possible role for PISD-LD in neutral lipid storage, we monitored incorporation of ^14^C oleate into triacylglcerol (TAG) in PISD and control RNAi HeLa cells (Fig. 6). Initially, we sought to eliminate PISD-LD expression using siRNAs directed against sequences within the unique segment of the mRNA, however, no effective siRNA could be identified, so we instead used an siRNA directed to the common region to reduce both PISD-LD and PISD-M. Using this approach, it was possible to reduce total PISD protein levels (determined by immunoblotting of cell lysates) by >90%. The gross appearance of LDs (number/cell, size), visualized by BODIPY staining and fluorescence microscopy, in these cells was indistinguishable from that of control cells. Next, PISD-depleted cells were incubated with 500 μM oleate (0.5 μCi of ^14^C oleate/6 well as tracer) for 1, 3, or 16 hours, lipids were extracted, separated by thin layer chromatography, and TAG was quantified. The results show that there is a substantial reduction (∼35% at the 16 hour time point) in the incorporation of oleate into TAG in PISD RNAi cells (Fig. 6). Thus, PISD promotes TAG storage.

**Figure 6:**
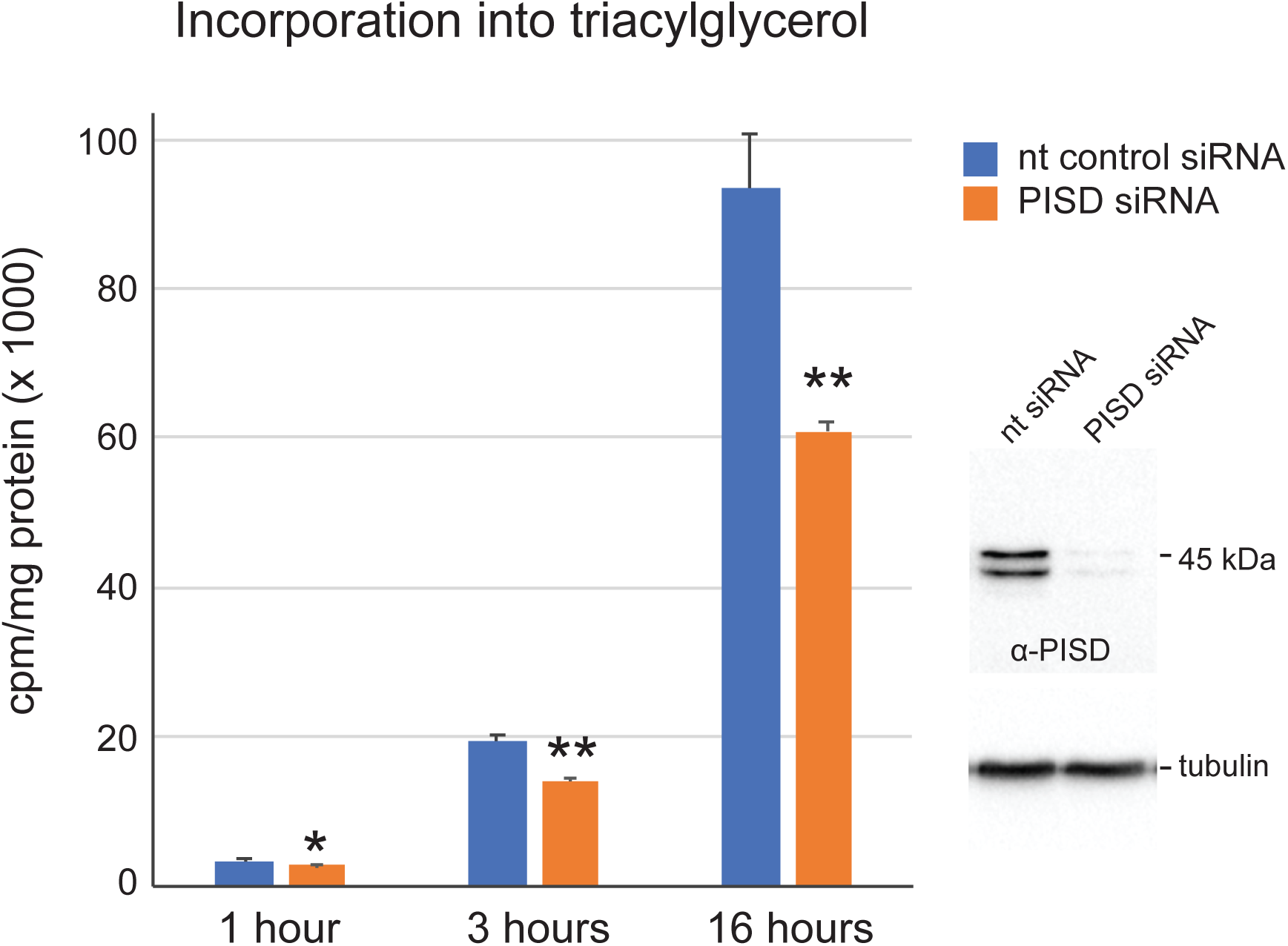
Decreased triacylglycerol synthesis by PISD RNAi cells. Forty eight hours after transfecting cells with siRNA directed against a sequence common to both PISD forms, the cell culture medium was supplemented with 500 μM oleate (0.5 μCi of ^14^C oleate/6 well as tracer) for 1, 3, or 16 hours. ^14^C oleate incorporation into TAG was measured over time and normalized to total cell protein. The mean (±s.d.) of triplicate measurements are plotted and statistical significance is indicated (** P < 0.01; * P < 0.05). Data are representative of two biological replicate experiments. To the right are shown immunoblots using antisera to PISD or tubulin of whole cell lysates prepared from control and PISD siRNA treated cells.

What is the physiological role of PISD in LD biology? LD biogenesis is initiated within the ER membrane, and proceeds via a maturation pathway resulting in the production of a mature LD containing a neutral lipid core and an ensemble of LD-associated proteins (Walther and Farese, 2012). A growing body of evidence indicates that interfacial surface tension of the LD monolayer controls association of LDs with ER membrane and LD size by influencing the recruitment, residence, and activities of enzymes that mediate neutral lipid storage and metabolism (Ben M’barek et al., 2017; Prevost et al., 2018). For example, the PE:PC ratio of the LD monolayer is critical for determining recruitment and activation of CCTα by influencing lipid packing defects that are sensed by an amphipathic alpha helix in CCTα (Arnold et al., 1997; Krahmer et al., 2011). It is notable that PS, the substrate of PSD, is largely absent from purified LDs, even though LDs are produced within the ER membrane, where PS is synthesized (Tauchi-Sato et al., 2002). We therefore speculate that the activity of PISD-LD on the surface of the LD ensures that PS, which may be incorporated into the LD from bulk ER membrane, does not accumulate on the surface of the LD. The effect of PISD depletion on TAG synthesis is not via deficient recruitment or activation of CCTα because depletion of CCTα results in an increase in TAG levels (Krahmer et al., 2011). However, many other LD residents, including acyltransferases involved in TAG synthesis (e.g., acyl-CoA:diacylglycerol acyltransferases) associate with the LD surface and it will be interesting to systematically examine targeting and activities of other LD-localized enzymes in cells deficient in PISD-LD. Our findings reveal a previously unknown physiological control point for cellular lipid metabolism linked to PE synthesis by PISD.

## Materials and Methods

### In silico analyses

Mitochondrion targeting sequences were predicated by MitoProt II (Claros and Vincens, 1996), secondary structures were predicated by Phyre2 (Kelley et al., 2015), and amphipathic helices were predicted by HELIQUEST (Gautier et al., 2008). Hydropathy plots were generated by tthe Kyte-Doolittle algorithm within the ProtScale program at http://web.expasy.org/protscale/.

### Molecular biology

Chemical reagents were purchased from Sigma-Aldrich (St. Louis, MO) unless otherwise indicated. RNA was isolated from Hela cells harvested from confluent cultures according to the manufacturer’s instructions using the RNeasy Mini Kit (Qiagen, Valencia, CA). RNA was used as a template for reverse-transcriptase polymerase chain reaction (RT-PCR) using the RevertAid First Strand cDNA Synthesis Kit (Thermo Fisher Scientific, USA). The pPISD-LD-eGFP vector was used as substrate to generate different mutants of PISD-LD by Q5® Site-Directed Mutagenesis kit (NEB). All final PCR-generated DNAs were confirmed by DNA sequencing. Oligonucleotide primers used for this study will be provided upon request.

### Cell fractionations

To prepare whole cell lysates, ∼2×10^6^ cells were harvested by scraping and washed twice with PBS. Cells were lysed in 150 µl 2x SDS-PAGE sample buffer. Lysates were boiled for 10 min at 95°C, centrifuged (13,000 *g*, 10 min, 4°C), and the supernatants transferred to new tubes.

Mitochondria were isolated from HeLa cells expressing PISD-LD-eGFP according to a published procedure (Parone et al., 2006) with the following modifications. Cells were lysed by 10 passages through a 25-gauge needle and the suspension was centrifuged at 500 *g* at 4°C for 5 min. The resulting supernatant was further centrifuged for 5 min at 10,000 *g* at 4°C to pellet the mitochondria. The pellet containing mitochondria was washed once with MB buffer [210 Mm mannitol, 70 mM sucrose, 10 mM HEPES, pH 7.5, 1 mM EDTA, 1X cOmplete protease inhibitor cocktail (Roche)] and then resuspended in MB buffer.

### SDS-PAGE and Western blotting

For immunoblotting, samples were resolved by 10% SDS-PAGE and transferred to 0.45 μm Nitrocellulose membrane (Bio-Rad). Blots were blocked and probed with mouse monoclonal anti-GFP (Sigma; 1:1000 dilution), mouse monoclonal anti-*β*-actin (Cell Signaling; 1:1000 dilution), rabbit anti-PISD (Santa Cruz Biotechnology; 1:1000 dilution), and rabbit anti-Tom20 (Santa Cruz Biotechnology; 1:400 dilution). Protein was visualized with either ECL Horseradish Peroxidase linked anti-rabbit or anti-mouse (Santa Cruz Biotechnology) secondary antibody (diluted 1:10,000) and SuperSignal® West Femto Maximum Sensitivity Substrate (Thermo Scientific, Rockford, IL).

### Cell Culture

HeLa T-REx cells (Invitrogen) were cultured in DMEM supplemented with 10% FBS (Gibco) and maintained in 5% CO_2_ at 37°C. Cells were transfected with Lipofectamine 2000 Transfection Reagent (Invitrogen) using 40-50 ng of plasmid DNA (unless otherwise indicated) mixed with 1µl of Lipofectamine 2000 and cultured for 16-20 hours prior to analysis.

For the lipid starvation experiment shown in Figure 2, transfected cells were treated with 100 µM oleic acid and 0.5 µM BODIPY Red (Invitrogen) for 24 hrs. Cells were washed three times with PBS and then incubated in media supplemented with 5% delipidated fetal bovine serum (Gemini BioProducts) for 0, 24 and 48 hrs.

### Live-cell microscopy and Image analysis

Image stacks of cells were collected at 0.3 μm z increments on a DeltaVision Elite workstation (Applied Precision) based on an inverted microscope (IX-70; Olympus) using a 100×, 1.4 NA or 60x, 1.4 NA oil immersion lens. Images were captured at 22°C with a sCMOS camera (CoolSnap HQ; Photometrics) and deconvolved with softWoRx version 6.0 using the iterative-constrained algorithm and the measured point spread function. Background signal was subtracted from images, which were then saved as JPEGs that were colored, denoised, and adjusted in brightness/ contrast/gamma with the program Fiji (Schindelin et al., 2012).

To localize GFP-tagged fusion proteins with BODIPY Red (Molecular Probes), transfected cells were grown overnight (16-20h) in culture media with 0.5 µM BODIPY Red. Cells were washed three times with PBS and medium was replaced with Live cell imaging solution (Molecular Probes). To localize GFP-tagged fusion proteins with MitoTracker Red (Molecular Probes), after 16h transfection, cells were rinsed three times with pre-warmed serum free media DMEM. Subsequently, cells were incubated with pre-warmed serum free media containing 25 nM MitoTracker Red for 10 min at 37°C. Cells were washed with PBS, medium was replaced with Live cell imaging solution and then imaged.

### [14C]-Oleic acid labelling of lipids, lipid extraction and thin layer chromatography

HeLa cells were transfected with Lipofectamine 2000 Transfection Reagent (Invitrogen). After 48 hours of transfection, cells were loaded with 500 μM oleate (0.5 μCi of ^14^C oleate/6 well as tracer) for 1, 3, or 16 hours. Cells were washed with phosphate buffer saline for 3 times. Lipids were extracted directly from 6-well cell-culture plates by adding hexane:isopropanol mixture (3:2) and gentle shaking for 10 min. Lipids were dried under nitrogen stream and separated by TLC using hexane:diethyl ether: acetic acid (80:20:1) solvent system. Thin layer chromatography (TLC) plates were exposed to a phosphor imaging cassette overnight and revealed by Typhoon FLA 7000 phosphor imager. Lipids on TLC plates were stained with iodine vapor; bands were scraped and quantified by liquid scintillation counter. After lipid extraction from 6-well plates, 400 μl of lysis buffer (0.3 N NaOH and 0.1% SDS) was added to each well and kept for shaking for 3 h to extract proteins from the cells. Total protein amount was measured by Bio-Rad *DC* Protein Assay kit.

### Statistical analyses

The student’s unpaired t test was used for statistical analyses using Prism (GraphPad software). Statistical significance is indicated as follows: ** *P* < 0.01; * *P* < 0.05.

## Acknowledgements

Research reported in this publication was supported by the National Institute of General Medical Sciences of the National Institutes of Health under award number GM060221 (to C.G.B.).C.C was supported by ADA mentor-based fellowship grant (7-12-MN-18 to C.C. and R.V.F), by the National Institute of National Institute of Diabetes and Digestive and Kidney Diseases of the National Institutes of Health under award number DK101579 (to T.C.W and R.V.F), the Mathers foundation (to T.C.W.). TCW is an investigator of the Howard Hughes Medical Institute.

## Competing interests

No competing interests declared.

## Abbreviations and nomenclature

CBB: coomassie brilliant blue
CCT: CTP:phosphocholine cytidylyltransferase
GAPDH: Glyceraldehyde 3-phosphate dehydrogenase
LD: lipid droplet
LFA: lipid free medium
OA: oleic acid
PE: phosphatidylethanolamine
PS: phosphatidylserine
PSD: phosphatidylserine decarboxylase
PISD1: phosphatidylserine decarboxylase proenzyme
PISD-LD: PISD isoform that localizes to the lipid droplet
PISD-M: PISD isoform that localizes solely to the mitochondrion
TAG: triacylglycerol

## Notes

### Competing Interest Statement

The authors have declared no competing interest.

